# Dysfunction of ventral tegmental area GABA neurons causes mania-like behavior

**DOI:** 10.1101/684142

**Authors:** Xiao Yu, Wei Ba, Guangchao Zhao, Ying Ma, Edward C. Harding, Lu Yin, Dan Wang, Youran Shi, Alexei L. Vyssotski, Hailong Dong, Nicholas P. Franks, William Wisden

## Abstract

The ventral tegmental area (VTA), an important source of dopamine, regulates goal- and reward-directed and social behaviors, wakefulness and sleep. Hyperactivation of dopamine neurons generates behavioral pathologies. But any roles of non-dopamine VTA neurons in psychiatric illness have been little explored. Lesioning or chemogenetically inhibiting VTA GABAergic (VTA*^Vgat^*) neurons generated persistent wakefulness with mania-like qualities: locomotor activity was increased; sensitivity to D-amphetamine was heightened; immobility times decreased on the tail suspension and forced swim tests; and sucrose preference increased. Furthermore, after sleep deprivation, mice with lesioned VTA*^Vgat^* neurons did not catch up on the lost NREM sleep, even though they were starting from an already highly sleep-deprived baseline, suggesting that the sleep homeostasis process was bypassed. The mania-like behaviors, including the sleep loss, were reversed by the mood-stabilizing drug valproate, and re-emerged when valproate treatment was stopped. Lithium salts, however, had no effect. The mania like-behaviors partially depended on dopamine, because giving D1/D2/D3 receptor antagonists partially restored the behaviors, but also on VTA*^Vgat^* projections to the lateral hypothalamus (LH). Optically or chemogenetically inhibiting VTA*^Vgat^* terminals in the LH elevated locomotion and decreased immobility time during the tail suspension and forced swimming tests. VTA*^Vgat^* neurons are centrally positioned to help set an animal’s (and human’s) level of mental and physical activity. Inputs that inhibit VTA*^Vgat^* neurons intensify wakefulness (increased activity, enhanced alertness and motivation), qualities useful for acute survival. Taken to the extreme, however, decreased or failed inhibition from VTA*^Vgat^* neurons produces mania-like qualities (hyperactivity, hedonia, decreased sleep).

## Introduction

During the mania phase of bipolar disorder, patients sleep little and have elevated mood (*e.g.* increased energy and hyperactivity, impulsivity, decreased depression)^1–5^. In mice, pathological hyperactivity and elevated mood can be generated by various gene mutations and deletions: *e.g.* ClockΔ19^6^, REV-erbα^7^, ErbB4 tyrosine kinase deletion in noradrenergic locus ceruleus cells^8^, GSK-3β overexpression^9^, dopamine transporter knockdown^10, 11^, SHANK2 knockout^12^, SHANK3 overexpression^13^, ANK3 disruptions^14^, ionotropic glutamate/AMPA receptor GluR1 knockout^15^, ionotropic glutamate/kainate GluR6 knockout^16^, phospholipase cγ1^8^, glutamate-cysteine ligase modifier unit knockout^17^, and the Na/K-ATPase α3 Myshkin (Myk/+) mutation ^18–20^. Some of these gene manipulations alter excitation-inhibition (E-I) balance^8, 13, 15, 16, 21^, and/or elevate catecholamines^6–8, 10, 22^, suggesting common themes that could underlie the emergence of some types of mania.

Both these themes come together in the ventral tegmental area (VTA). The VTA an important source of dopamine, regulates goal- and reward-directed and social behaviors ^23, 24^, as well as wakefulness and sleep ^25–27^. Exciting VTA dopamine neurons with well-chosen rhythms can produce mania-like behaviors in the day and euthymia at night ^28^. In addition to dopamine neurons, there is a rich heterogeneity of glutamate and GABA neurons in the different anatomical subdomains of the VTA, and neurotransmitter co-release (e.g. dopamine - glutamate, GABA-glutamate, GABA-dopamine) from VTA neurons is common^23, 29–31^.

For this paper, we focus on the midline VTA, which contains GABA (VTA*^Vgat^*) and glutamate/nitric oxide synthase (VTA*^Vglut^*^2^) neurons^23, 32, 33^. These VTA*^Vgat^* and VTA*^Vglut2/NOS1^* neurons inhibit and excite, respectively, the dopamine cells, but also, by projecting out of the VTA, exert effects independent of dopamine ^23^. The VTA*^Vgat^* neurons co-release GABA and glutamate^31^, but the majority of these VTA neurons’ actions locally are GABAergic^26^. Locally they mostly inhibit VTA*^Vglut^*^2^, VTA^DA^ and VTA*^Vglut2^*^/DA^ cells, but also elicit a small number of pure excitatory responses ^26^. Chemogenetic inhibition and chronic lesion of midline VTA*^Vgat^* neurons causes sustained wakefulness^26, 27^, and excitation of VTA*^Vglut2^* cells and VTA dopamine cells also causes wakefulness^26 25^. VTA*^Vgat^* neurons limit arousal by inhibiting dopamine neurons and via projections to the lateral hypothalamus (LH)^26^; VTA*^Vglut2^* neurons produce wakefulness, also independently of dopamine, by projecting to the LH and nucleus accumbens ^26^.

Here, we characterize the type of wakefulness produced by inhibiting or lesioning the VTA*^Vgat^* neurons or exciting VTA*^Vglut2^* neurons. We find that the wakefulness induced by diminishing or removing VTA*^Vgat^* neuron inhibition contains behavioral endophenotypes that are mania-like, and are treatable with valproate, although not with lithium. On the other hand, the extended but quiet wakefulness produced by activating midline VTA*^Vglut2^* neurons has no endophenotypes characteristic of mania. We suggest that the mania-like symptoms resulting from diminished VTA*^Vgat^* function are generated by changing the E-I balance in both the VTA and the LH.

## Material & Methods

### Mice

All experiments were performed in accordance with the UK Home Office Animal Procedures Act (1986); all procedures were approved by the Imperial College Ethical Review Committee and the Ethics Committee for Animal Experimentation of Xijing Hospital, Xi’an, and were conducted according to the Guidelines for Animal Experimentation of the Chinese Council institutes. The following strains of mice were used: *Vgat-ires-Cre: Slc32a1^tm2(cre)Lowl^*/*J* kindly provided by B.B. Lowell, JAX stock 016962 ^34^; *Vglut2-ires-Cre: Slc17a6^tm2(cre)Lowl^/J*, kindly provided by B.B. Lowell, JAX stock 016963 ^34^. Mice were maintained on a 12 hr:12 hr light:dark cycle at constant temperature and humidity with *ad libitum* food and water.

### AAV

*pAAV-hSyn-DIO-hM4Di-mCherry*, *pAAV-hSyn-DIO-hM3Dq-mCherry* and *pAAV-hSyn-DIO-mCherry* were gifts from Bryan L. Roth (Addgene plasmid 44362, 44361 and 50459) ^35^; *pAAV-EF1α-DIO-taCASP3-TEV* was a gift from Nirao Shah (Addgene plasmid 45580) ^36^; we packaged the transgenes into AAV capsids (capsid serotype 1/2) in house as described previously ^37, 38^. *rAAV2/9-EF1a-DIO-eNpHR3.0-mCherry* was packaged by BrainVTA (Wuhan, China).

### Surgery

Mice were anesthetized with 2% isoflurane in oxygen by inhalation and received buprenorphine (0.1 mg/kg) and carprofen (5 mg/kg) injections, and then placed on a stereotaxic frame (Angle Two, Leica Microsystems, Milton Keynes, Buckinghamshire, UK). The AAV was injected through a stainless steel 33-gauge/15mm/PST3 internal cannula (Hamilton) attached to a 10 µl Hamilton syringe, at a rate of 0.1 µl min^−1^. For the AAV injections, virus was bilaterally injected into the VTA, 50 nl for each side of the VTA. The injection co-ordinates were VTA: (ML = ±0.35 mm, AP = −3.52 mm, DV = −4.25 mm). After injection, the cannula was left at the injection site for 5 min and then slowly pulled out. After injections, mice that were to undergo the sleep experiments were implanted with 3 gold-plated miniature screw electrodes (–1.5 mm Bregma, +1.5 mm midline; +1.5 mm Bregma, –1.5 mm midline; –1 mm Lambda, 0 mm midline – reference electrode) with two EMG wire (AS634, Cooner Wire, CA). The EMG electrodes were inserted between the neck musculature. The EEG-EMG device was affixed to the skull with Orthodontic Resin power and Orthodontic Resin liquid (Tocdental, UK). For the fiber optogenetic and chemogenetic experiments, after virus injection above the VTA (ML = ±0.35 mm; AP = −3.52 mm; DV = −4.25 mm), mice received surgical bilateral implantations above the LH of a monofiberoptic cannula (ML = ±0.36 mm; AP = −3.54 mm; DV = −4.0 mm) (200 µm; Doric Lenses, Inc., Quebec, Canada) or guide cannula (ML = ±0.36 mm; AP = −3.54 mm; DV = −3.5 mm) (World precision instruments, USA) for CNO delivery.

### Drug treatments

For all chemogenetic and optogenetic experiments, mice were split into random groups that received either saline or CNO injection or with or without opto-stimulation. For dopamine antagonists injection, mice were split into random groups that received either vehicle or antagonists injection. For acute treatments with lithium or valproate, mice were received vehicle injection and behaviors (locomotion and tail-suspension) were tested, and after 1-2 week, same group of mice were received lithium or valproate injection. For acute valproate treatment with sleep experiment, mice were split into two random groups that received vehicle injection or valproate injection, and 24 hour sleep were recorded. For the Li-H_2_O treatment, mice were first placed with normal water, and behaviors were tested, and same group of mice were placed with Li-H_2_O for 2 weeks.

### Chemogenetics

Clozapine-N-oxide (CNO) (C0832, Sigma-Aldrich, dissolved in saline, 1 mg/kg) or saline was injected *i.p*. 30 minutes before the start of the behavioral tests. For VTA*^Vgat^*-hM4Di mice or VTA*^Vglut2^*-hM3Dq mice, CNO or saline was injected during the “lights off” active phase. 1 μl CNO (5 mg/kg) was infused into the LH through the guide cannula at a rate of 0.1 µl min^−1^ with an injector needle (33-gauge, Hamilton)

### D-amphetamine

D-Amphetamine (2813/100, Tocris Bioscience, dissolved in saline, 2 mg/kg) or saline was injected *i.p*. into the mice ^13, 19^, and the mice were assessed directly after D-amphetamine injection.

### Valproate and lithium

Sodium valproate (2815, TOCRIS, dissolved in saline, 200 mg/kg), LiCl (Sigma-Aldrich, dissolved in saline, 100 mg/kg) or saline (vehicle) was injected *i.p*. into the mice. Valproate or LiCl treatments were as previously reported ^13, 39^: valproate or LiCl were injected 3 times (10:00, 14:00 and 17:00) one day before the behavioral assays; and during the day of the behavioral assay, valproate or LiCl was injected 2 times (10:00, 14:00). Locomotion or tail-suspension tests were performed 30 min after injection (14:30). After the locomotion or tail-suspension tests (see below), mice received a final valproate injection (17:00), and then the 24 hour sleep-wake recordings were then performed (see section “**EEG analysis, sleep-wake behavior and sleep deprivation**”). For the Li-H_2_O treatment, mice were treated with LiCl in drinking water (300 mg/L) for 2 weeks.

### Dopamine receptor antagonists

Dopamine antagonists SCH-23390 (0.03 mg/kg, dissolved in saline) and raclopride (1 mg/kg, dissolved in saline), for D1 and D2/D3 receptors respectively, were injected serially *i.p.* 30 minutes before the behavioral tests.

### Timing of behavioral tests

All behavioral tests (see below) took place during the “lights off” phase of the light-dark cycle, when the mice were most active, except for the sleep deprivation experiments (see section “**EEG analysis, sleep-wake behavior and sleep deprivation”**).

### Locomotor activity

For chemogenetic experiments, the mice were received saline or CNO injection, the locomotor activity was detected 30 min after injection, and 1-2 week later, the same mice were received CNO or saline injection, and the locomotor activity was detected 30 min after injection. For D-Amphetamine experiments, mice were first put into the open field for 30 min, and the mice were received vehicle or amphetamine injection. The locomotor activity was detected straight after injection for 1 hour. The locomotor activity was detected in an activity test chamber (Med Associates, Inc) with an ANY-maze video tracking system (FUJIFILM co.), and measured by ANY-maze software (Stoelting Co. US.).

### Home cage activity

Mice were habituated in the cage for 24 h, and the activities were recorded by video tracking system for 1 h, and measured by ANY-maze software (Stoelting Co. US.).

### Tail-suspension test (TST)

The TST was performed as described^13^. After a habituation period in the test room, mice were suspended 60 cm above the floor by their tails by taping the tail tip. Behaviors were video recorded and blindly scored manually and measured by ANY-maze software (Stoelting Co. US.).

### Forced swimming test (FST)

Mice were placed in a borosilicate glass cylinder (5L, 18 cm diameter, 27 cm high) filled with water (25 °C, water depth 14 cm) for 6 min. The immobility time during the last 4 min was manually measured. Immobility time was defined as the time spent without any movements except for a single limb paddling to maintain flotation.

### Sucrose preference test (SPT)

Mice were habituated to a dual delivery system (one bottle with water and one bottle with 1% sucrose) for 3 days. Sucrose preference was then assessed over 3 consecutive days. For the chemogenetic experiments, animals were water-restricted overnight before saline or CNO injection. 30 min after saline or CNO injection, the mice were given free access to the 2-water delivery system for 4 hours. Sucrose preference (%) was calculated as (weight of sucrose consumed) / (weight of water consumed + weight of sucrose consumed) × 100%.

### EEG analysis, sleep-wake behavior and sleep deprivation

EEG and EMG signals were recorded using Neurologger 2A devices^26, 40^. NREM sleep and wake states were automatically classified using a sleep analysis software Spike2 and manually scored. The sleep deprivation protocol was as described previously^41^. At the start of ‘lights on’ sleep period, mice fitted with Neurologgers were put into novel cages, and at one-hour intervals, novel objects were introduced. After 5 hours sleep deprivation, mice were then put back into their home cages for 19 hours. In total, 24 hour sleep-wake state were recorded. For the sleep experiments with dopamine receptor antagonists, mice were given SCH-23390 (0.03 mg/kg) and raclopride (2 mg/kg) injections in the middle of the ‘lights off’ active period.

### Optogenetic stimulation

For the optogenetic behavioral experiments, a fiber patch cord was connected to the laser generator, and dual optic fibers were connected to the fiber patch through a rotary joint (ThinkerTech Nanjing BioScience Inc. China.). Before the experiments, a monofiberoptic cannula was connected to the fiber patch cord. VTA*^Vgat^*-eNpHR-mCherry*→* LH mice were bilaterally opto-stimulated (20 Hz, 0.5 s duration with 0.5 s interval, 593 nm, 20 μW) in the LH during the “lights off” phase. The stimulation was given for the duration of the behavioral tests.

### Immunohistochemistry

These procedures were carried as described previously^26^. Primary antibodies used were: rat monoclonal mCherry (1:2000, Clontech); mouse monoclonal tyrosine hydroxylase (TH) (1:1000, Molecular Probes); secondary antibodies were Alexa Fluor 488 goat anti-mouse, Alexa Fluor 594 goat anti-rat (1:1000, Invitrogen Molecular Probes, UK). Slices were mounted on slides, embedded in Mowiol (with 4,6-diamidino-2-phenylindole), cover-slipped, and analyzed using an upright fluorescent microscope (Nikon Eclipse 80i, Nikon).

## Results

### Chemogenetic inhibition or lesioning of VTA*^Vgat^* neurons produces increased locomotor activity and sensitivity to amphetamine

We delivered AAV-DIO-hM4Di-mCherry into the VTA of *Vgat-ires-cre* mice to express hM4Di-mCherry (an inhibitory receptor activated by the CNO ligand) specifically in VTA*^Vgat^* neurons to generate VTA*^Vgat^*-hM4Di mice (Figure 1a)^26^. As previously^26^, we also ablated VTA*^Vgat^* neurons by injecting AAV-DIO-CASP3 and/or AAV-DIO-mCherry (as a control) into the VTA of *Vgat-ires-cre* mice to generate VTA*^Vgat^*-CASP3 mice and VTA*^Vgat^*-mCherry mice respectively. After 5-6 weeks post-injection, mCherry-positive VTA*^Vgat^* neurons were detected in control mice that had only received AAV-DIO-mCherry injections, whereas mCherry-positive VTA*^Vgat^* neurons were mostly absent in caspase-injected mice (Figure 1b), but tyrosine hydroxylase-positive (dopamine) neurons were still present (Figure 1b).

**Figure 1.**
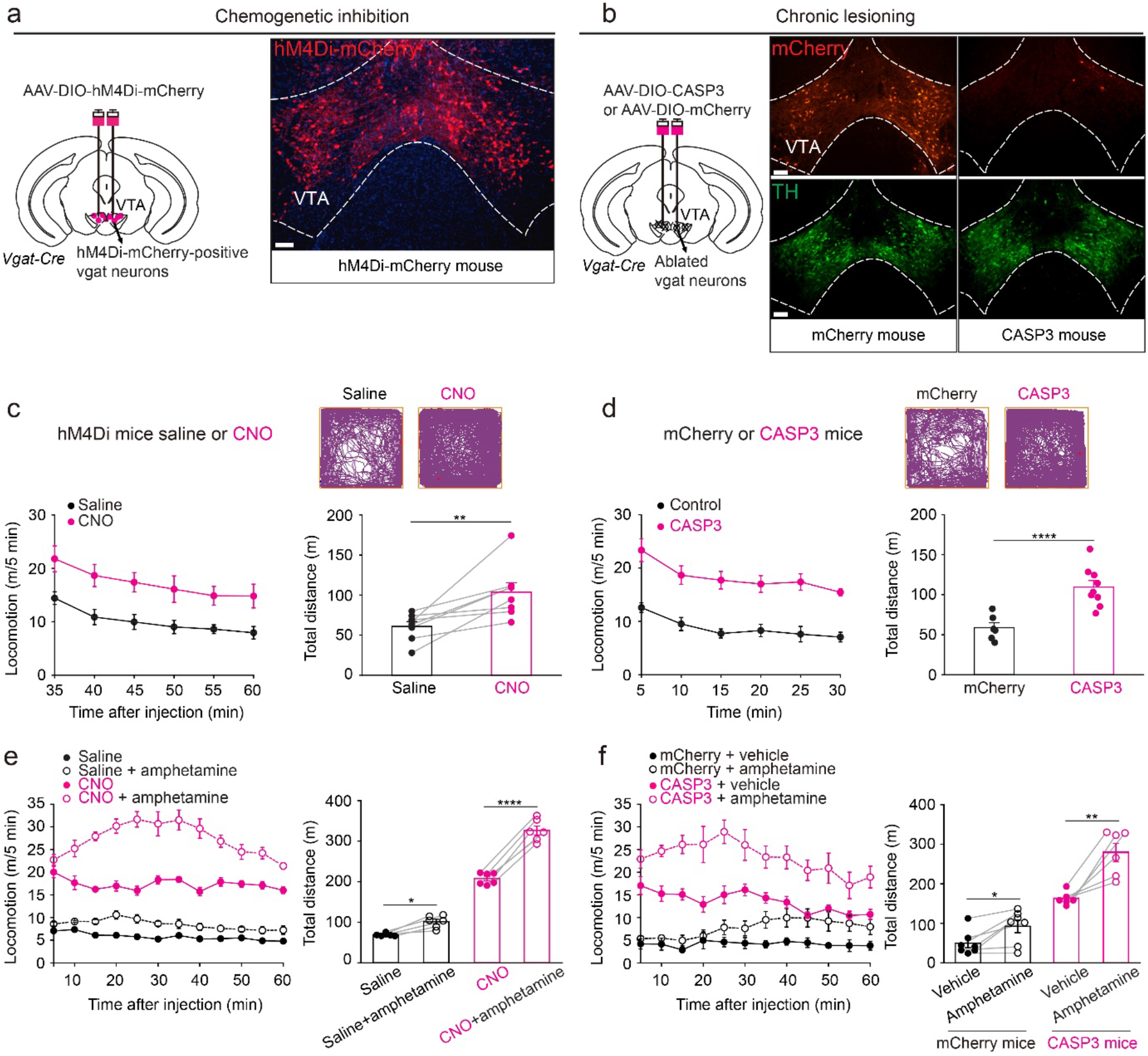
Selective inhibition or lesioning of VTA*^Vgat^* neurons increased locomotor activity and sensitivity to D-amphetamine-induced hyperlocomotion. (a) Generation of VTA*^Vgat^*-hM4Di mice. Staining by immunohistochemistry (mCherry, red) for hM4Di-mCherry expression in the VTA. Scale bar: 150 μm. (b) Generation of VTA*^Vgat^*-CASP3 and VTA*^Vgat^*-mCherry control mice. Immunohistochemical staining for mCherry (red) and dopamine neurons (TH, tyrosine hydroxylase, green) in the VTA of control mice that expressed mCherry in VTA^Vgat^ neurons (left-hand images), or in mice where VTA^Vgat^ neurons were lesioned with caspase (right-hand images). Scale bar: 150 μm. (c) Video-tracked paths, locomotion speed and distance travelled of VTA*^Vgat^*-hM4Di mice (n=8 mice) following CNO (1 mg/kg) injection compared with saline injection in the open field over a 30 minute period. Paired t-test, t(7)=−3.68, **p=0.007. (d) Video-tracked paths, locomotion speed and distance travelled of VTA*^Vgat^*-mCherry (n=8 mice) and VTA*^Vgat^*-CASP3 mice (n=9 mice) in the open field over a 30 minute period. Unpaired t-test, t(15)=−5.43, ****p=0.00006. (e) Locomotion speed and distance traveled of VTA*^Vgat^*-hM4Di mice (n=6 mice) in the open field after saline or D-amphetamine injection (subsequent to saline or CNO injection). Repeated measures two-way ANOVA and Bonferroni-Holm *post hoc* test. F(1,5)=66. Saline vs. saline + amphetamine t(5)=3.69, *p=0.01; CNO vs. CNO + amphetamine t(5)=13.6, ****p=0.000038. (f) Locomotion speed and distance traveled in the open field test of VTA*^Vgat^*-mCherry (n=7 mice) or VTA*^Vgat^*-CASP3 mice (n=6 mice) after vehicle or D-amphetamine injection. Two-way ANOVA and Bonferroni-Holm *post hoc* test. F_(mCherry or CASP3)_=93; F_(saline or amphetamine)_=26; F_(interaction)_=5.64; mCherry + vehicle vs. mCherry + amphetamine t(6)=−3.39, *p=0.01; CASP3 + vehicle vs. CASP3 + amphetamine t(5)=−4.97, **p=0.004. All error bars represent the SEM.

We next assessed locomotion behaviors of these animals. After CNO injection, VTA*^Vgat^*-hM4Di mice had higher locomotor activity in an open-field arena with more distance travelled (Figure 1c), and had elevated average and maximum speeds over 30 min compared with saline-injected VTA*^Vgat^*-hM4Di mice (Supplementary Figure 1a). Consistent with these chemogenetic inhibition results, VTA*^Vgat^*-CASP3 mice also produced hyperlocomotion in the open field – more distance travelled (Figure 1d), and higher average and maximum speeds over 30 min compared with the distance travelled by VTA*^Vgat^*-mCherry control mice (Figure 1d; Supplementary Figure 1b). VTA*^Vgat^*-CASP3 mice were also hyperactive in their home cages (Supplementary Figure 1c). Notably, the hyperlocomotion of the VTA*^Vgat^*-CASP3 mice was long-term, as seen when the mice were retested in the open field four months post lesion (Supplementary Figure 1d).

Mice suggested to have mania-like characteristics, or patients with bipolar-related mania, are hypersensitive to D-amphetamine^13, 19^, whereas mice and patients posited to have an attention deficit hyperactivity (ADHD)-like disorder become less active with D-amphetamine^42, 43^. Treating CNO-injected VTA*^Vgat^*-hM4Di mice and VTA*^Vgat^*-CASP3 mice with D-amphetamine significantly increased their locomotion speed above their high baseline speed, and increased the distance travelled (Figure 1e, f), suggesting that both groups of mice (VTA*^Vgat^*-CASP3 and CNO-injected VTA*^Vgat^*-hM4Di mice) were not ADHD-like.

### Acute inhibition or chronic lesioning of VTA*^Vgat^* neurons elevates mood

We assessed mood-related behaviors using the tail-suspension test (TST), the forced swimming test (FST), and the sucrose preference test (SPT). During the TST, the immobility times of CNO-injected VTA*^Vgat^*-hM4Di mice and VTA*^Vgat^*-CASP3 mice were greatly decreased compared with saline-injected VTA*^Vgat^*-hM4Di control mice and VTA*^Vgat^*-mCherry (Figure 2a, b). Both CNO-injected VTA*^Vgat^*-hM4Di mice and VTA*^Vgat^*-CASP3 mice also had decreased immobility times during the FST compared with saline-injected or VTA*^Vgat^*-mCherry controls (Figure 2c, d). This could suggest that reducing VTA*^Vgat^* neuronal output, by either chemogenetic inhibition or lesioning, produces less depressive-like behaviors. In addition, during the SPT, CNO-injected VTA*^Vgat^*-hM4Di mice and VTA*^Vgat^*-CASP3 mice consumed more sucrose than their control littermates (Figure 2e, f), possibly indicating a raised hedonic state.

**Figure 2.**
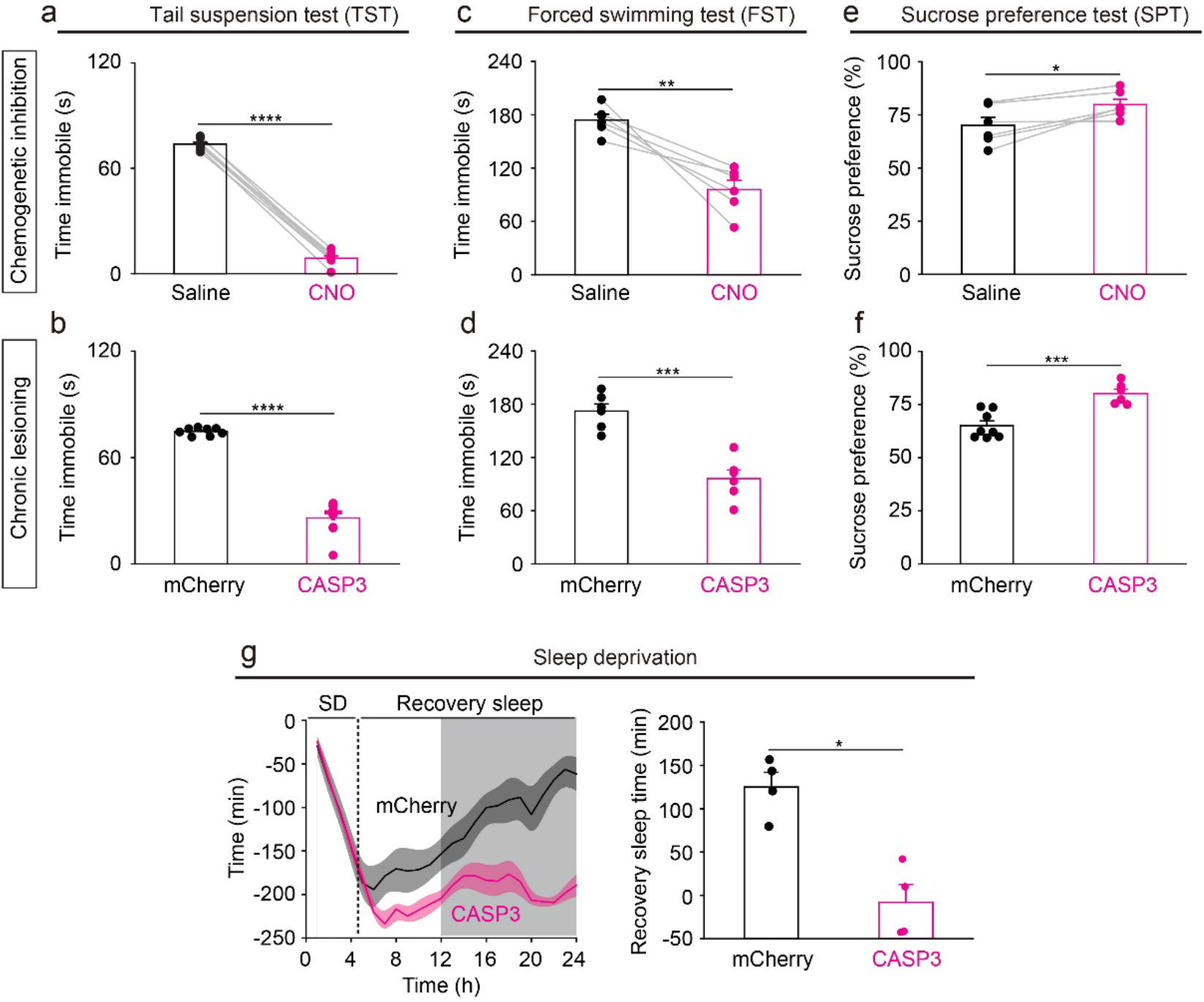
Dysfunction of VTA*^Vgat^* neurons elevates mood and impairs sleep homeostasis. (a) Immobility time during the tail suspension test (TST) of VTA*^Vgat^*-hM4Di mice (n=8 mice) after saline or CNO injection. Paired t-test, t(7)=53.8, ****p=1.98e^-10^. (b) Immobility time during the during TST of VTA*^Vgat^*-CASP3 (n=9 mice) or VTA*^Vgat^*-mCherry mice (n=8 mice). Unpaired t-test, t(15)=14.3, ****p=3.7e^-10^. (c) Immobility time during the forced swimming test (FST) of VTA*^Vgat^*-hM4Di mice (n=6 mice) after saline or CNO injection. Paired t-test, t(5)=5, **p=0.003. (d) Immobility time during the FST of VTA*^Vgat^*-CASP3 (n=6 mice) or VTA*^Vgat^*-mCherry mice (n=6 mice). Unpaired t-test, t(10)=6, ***p=0.0001. (e) Sucrose preference of VTA*^Vgat^*-hM4Di mice (n=6 mice) after saline or CNO injection. Paired t-test, t(5)=−3.5, *p=0.01. (f) Sucrose preference of VTA*^Vgat^*-CASP3 (n=6 mice) or VTA*^Vgat^*-mCherry mice (n=8 mice). Unpaired t-test, t(12)=−4.8, ***p=0.0004. (g) The accumulative recovery sleep (time in NREM sleep) after 5 hours of sleep deprivation (SD) of VTA*^Vgat^*-CASP3 (n=4 mice) or VTA*^Vgat^*-mCherry mice (n=4 mice). Mann-Whitney test, *p=0.03. The light and dark shading represents the “lights on” and “lights off” phases.

### Mice with lesioned VTA^Vgat^ neurons have less sleep need after sleep deprivation

VTA*^Vgat^*-CASP3 mice have a long-term sleep deficit^26^. To further characterize the need of VTA*^Vgat^*-CASP3 mice for sleep, we performed a sleep deprivation experiment. After sleep deprivation, there is usually a rebound in lost NREM sleep, a process termed sleep homeostasis^44^. Mice were kept awake for 5 hours with novel objects presented each hour, and were then tested to see if they caught up on lost sleep in their home cage. Control VTA*^Vgat^*-mCherry mice had rebound NREM sleep after 5 hours sleep deprivation, and thereby after 19 hours they had regained 90% of their sleep loss (Figure 2g). Surprisingly, the VTA*^Vgat^*-CASP3 mice, despite already starting from a chronic sleep-deprived baseline, did not catch up on lost NREM sleep after 5 hours continuous sleep deprivation (Figure 2g).

### The mania-like behavior produced by lesioning VTA^Vgat^ neurons can be pharmacologically rescued by valproate but not lithium

We assessed if lithium could treat the manic-like state of VTA*^Vgat^*-CASP3 mice (Supplementary Figure 2). However, neither acute (100 mg/kg) (Supplementary Figure 2a-e) nor chronic (300 mg/L) treatments (Supplementary Figure 2f-j) had any effects (Of note, chronic Li-H_2_O treatment decreased the immobility time of control mice during the TST (Supplementary Figure 2i)). We next examined valproate (200 mg/kg) treatments (Figure 3a). Acute injection of valproate did not affect the locomotor activity of VTA*^Vgat^*-mCherry control mice (Figure 3b). By contrast, the hyperlocomotor activity of VTA*^Vgat^*-CASP3 mice in the open field was restored down to control levels by valproate treatment (Figure 3c). However, 2 weeks after the acute valproate treatment was withdrawal from VTA*^Vgat^*-CASP3 mice, their hyperactivity in the open field had returned (Figure 3c). Similarly, during the tail suspension test, valproate treatment did not affect VTA*^Vgat^*-mCherry control mice (Figure 3d), but significantly increased the immobility time of VTA*^Vgat^*-CASP3 mice back up to control levels (Figure 3e). Two weeks after valproate had been removed, however, the abnormally high agitation of VTA*^Vgat^*-CASP3 mice had re-emerged (Figure 3e).

**Figure 3.**
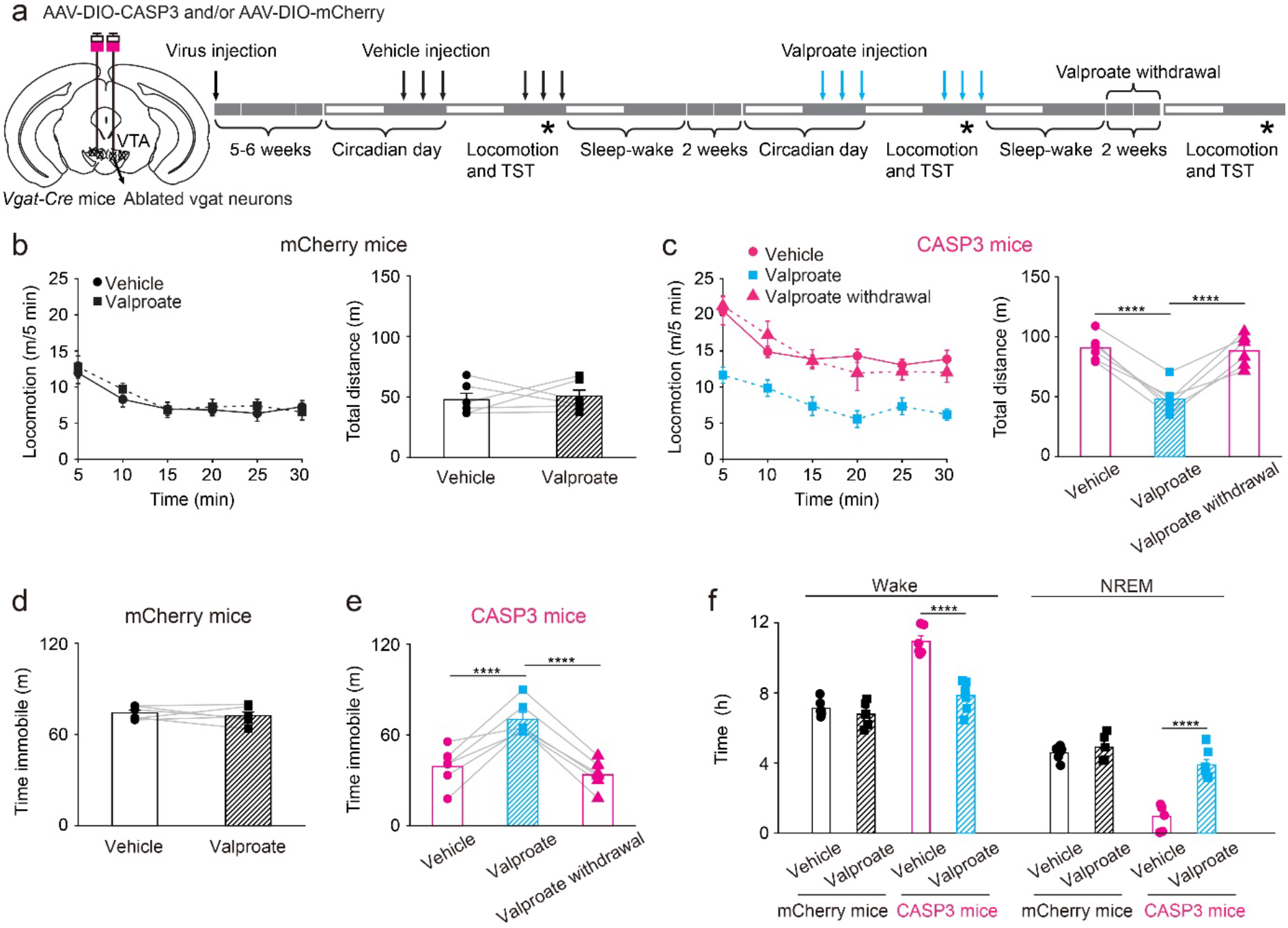
Valproic acid can successfully treat the mania-like behavior of VTA*^Vgat^*-CASP3 mice. (a) Pharmacological treatment protocol for valproic acid. The top arrows indicate vehicle or valproic acid injection (depending on mouse group) and the stars indicate when the behavioral experiments were undertaken. (b) Locomotion speed and distance travelled for VTA*^Vgat^*-mCherry mice (n=6 mice) that received either vehicle or valproic acid treatment. Paired t-test, t(5)=−0.36, p=0.73. (c) Locomotion speed and distance travelled for VTA*^Vgat^*-CASP3 mice (n=6 mice) received either vehicle or valproic acid treatment, or where the valproic acid treatment had been removed for 2 weeks. Repeated measures one-way ANOVA and Bonferroni-Holm post hoc test. F(2,10)=47; vehicle vs. valproate t(10)=8.68, ****p=0.000005; valproate vs. 2 weeks after valproate withdrawal t(10)=8.19, ****p=0.000009; vehicle vs. after valproate withdrawal t(10)=0.48, p=0.63. (d) Time spent immobile on the TST of VTA*^Vgat^*-mCherry mice (n=6 mice) received vehicle or valproate injection. Paired t-test, t(5)=0.71, p=0.5. (e) Time spent immobile on the TST of VTA*^Vgat^*-CASP3 mice (n=6 mice) received vehicle, valproate injection, or 2 weeks after valproate withdrawal. Repeated measures one-way ANOVA and Bonferroni-Holm post hoc test. F(2,10)=32; vehicle vs. valproate t(10)=7.41, ****p=0.00002; 2 weeks after valproate withdrawal; t(10)=6.32, ****p=0.00008; vehicle vs. remove t(10)=1.08, p=0.3. (f) Percentage and time of wake and NREM for VTA*^Vgat^*-mCherry mice and VTA*^Vgat^*- CASP3 mice over the 12 hours “lights off” period that received saline or valproate injection. mCherry + vehicle n=6 mice; mCherry + valproate n=5 mice; CASP3 + vehicle n=6 mice; CASP3 + valproate n=7 mice. Two-way ANOVA and Bonferroni-Holm *post hoc* test. Wake: F_(mCherry or CASP3)_=66; F_(Vehicle or valproate)_=17; F_(interaction)_=21;mCherry + vehicle vs. mCherry + valproate t(9)=0.38, p=0.38; CASP3 + vehicle vs. CASP3 + valproate t(11)=6.89, ****p=0.00002. NREM: F_(mCherry or CASP3)_=65; F_(Vehicle or valproate)_=32; F_(interaction)_=20; mCherry + vehicle vs. mCherry + valproate t(9)=−0.92, p=0.37; CASP3 + vehicle vs. CASP3 + valproate t(11)=−6.85, ****p=0.00002. All error bars represent the SEM.

We assessed if valproate could reduce the sustained wakefulness of VTA*^Vgat^*-CASP3 mice. The wake time of VTA*^Vgat^*-CASP3 mice significantly decreased after treatment with valproate, whereas NREM sleep time substantially increased (Figure 3f). VTA*^Vgat^*-CASP3 mice also have a pathological sleep architecture with fewer episodes of wake and NREM sleep, prolonged duration of each wake episode, and substantially decreased numbers of wake-NREM transitions^26^. Valproate treatment normalized the sleep-wake architecture of the VTA*^Vgat^*-CASP3 mice: the episode number (Supplementary Figure 3a), episode duration of wake were restored to control levels (Supplementary Figure 3b), as were the number of transitions between wake and NREM sleep (Supplementary Figure 3c).

### The extended wakefulness produced by activating VTA*^Vglut2^* neurons is not mania-like

Activating VTA*^Vglut2^* neurons also promotes wakefulness^26^. As an internal control to eliminate the possibility that the form of artificially-induced wake in these mouse models is likely to be mania-like, we expressed the excitatory chemogenetic hM3Dq receptor in VTA*^Vglut2^* neurons (VTA*^Vglut2^*-hM3Dq mice; Figure 4a)^8^. Chemogenetic activation of VTA*^Vglut2^* neurons by CNO injection did not alter the locomotor activity or distance travelled of VTA*^Vglut2^*-hM3Dq mice compared with saline injected mice (Figure 4b) (see also data in ref.^26^). Moreover, the immobility time during the TST and FST did not differ between CNO injected- and saline injected VTA*^Vglut2^*-hM3Dq mice (Figure 4c, d). Thus, the activation of VTA*^Vglut2^* neurons did not produce hyperactivity and mood-related deficits, suggesting a different kind of wakefulness from that generated by inhibition of VTA*^Vgat^* neurons.

**Figure 4.**
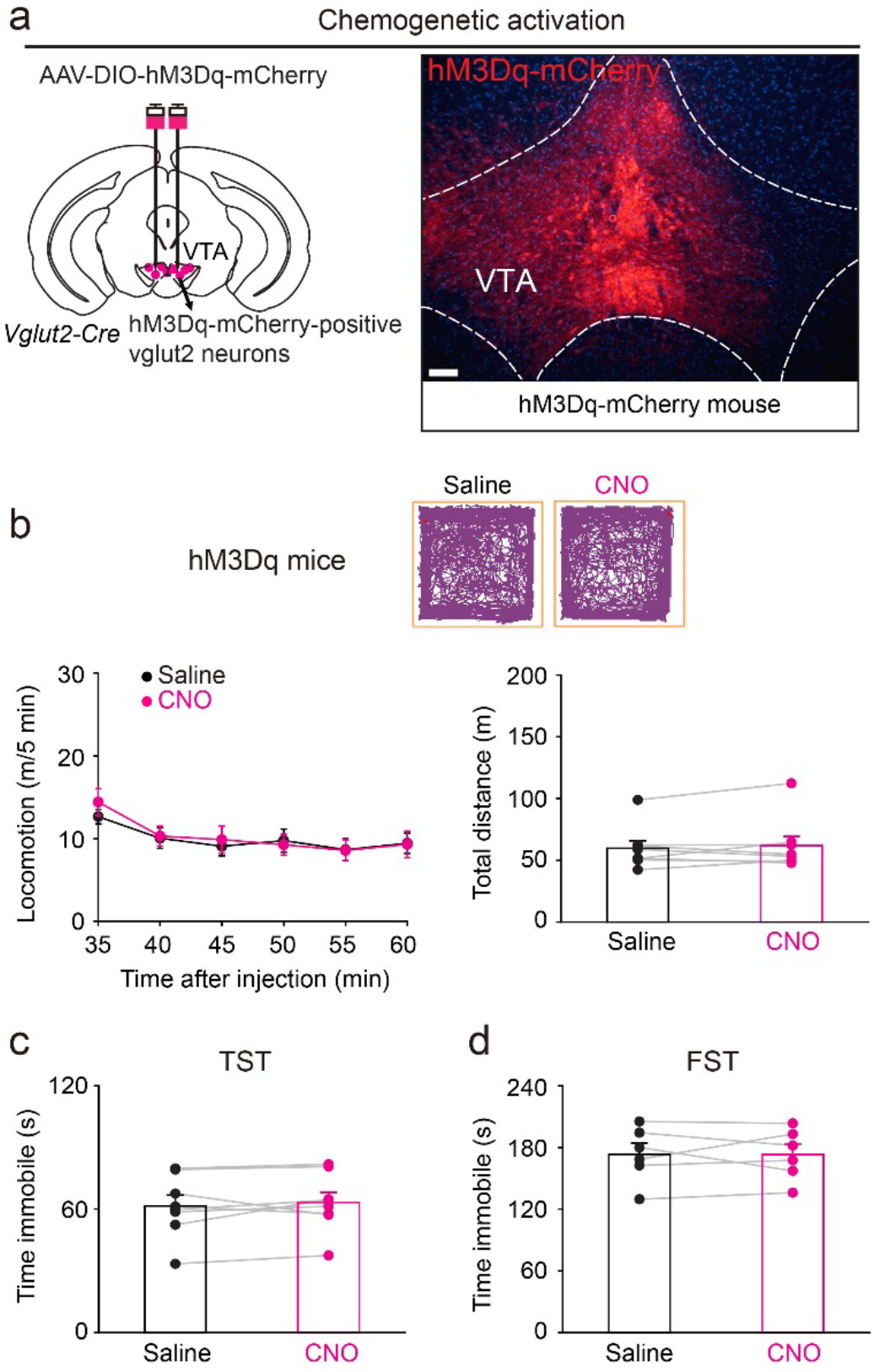
Activation of VTA*^Vglut2^* neurons does not produce mania-like behaviors. (a) Generation of VTA*^Vglut2^*-hM3Dq mice. Staining by immunohistochemistry (mCherry) for hM3Dq-mCherry expression in the VTA. Scale bar: 150 μm. (b) Video-tracked paths, locomotion speed and distance travelled of VTA*^Vglut2^*-hM3Dq mice (n=8 mice) following CNO (1 mg/kg) injection compared with saline injection in the open field arena over a 30 minute period. Paired t-test, t(7)=−0.72, p=0.49. (c) Immobility time during the tail suspension test (TST) of VTA*^Vglut2^*-hM3Dq mice (n=8 mice) after saline or CNO injection. Paired t-test, t(7)=−0.73, p=0.88 (d) Immobility time during the forced swimming test (FST) of VTA*^Vglut2^*-hM3Dq mice (n=6 mice) after saline or CNO injection. Paired t-test, t(4)=−0.01, p=0.98.

### VTA GABAergic neurons contribute to mania-like behaviors via dopamine signaling and projections to the LH

To examine whether the hyperactive behaviors in the VTA*^Vgat^*-CASP3 mice were produced by increased dopamine signaling, we gave a dopamine receptor antagonist mixture *i.p.* (containing SCH-23390 and raclopride for D1 and D2/D3 receptors, respectively). In the open-field test, the dopamine antagonists reduced the hyperlocomotion and distance travelled of VTA*^Vgat^*-mCherry control mice (Figure 5a), but only had a subtle effect on VTA*^Vgat^*-CASP3 mice (Figure 5a). In the TST, dopamine receptor antagonists increased the immobility time of both VTA*^Vgat^*-mCherry control mice and VTA*^Vgat^*-CASP3 mice (Figure 5b). However, the immobility time of VTA*^Vgat^*-CASP3 mice that received dopamine receptor antagonist injection was still significant lower than VTA*^Vgat^*-mCherry injected with dopamine receptor antagonists (Figure 5b). We also found that the extended wakefulness of VTA*^Vgat^*-CASP3 mice was substantially attenuated by the dopamine receptor antagonists (Supplementary Figure 4). The above results suggest that blocking dopamine signaling partially reduces the mania-like behaviors of VTA*^Vgat^*-CASP3 mice.

**Figure 5.**
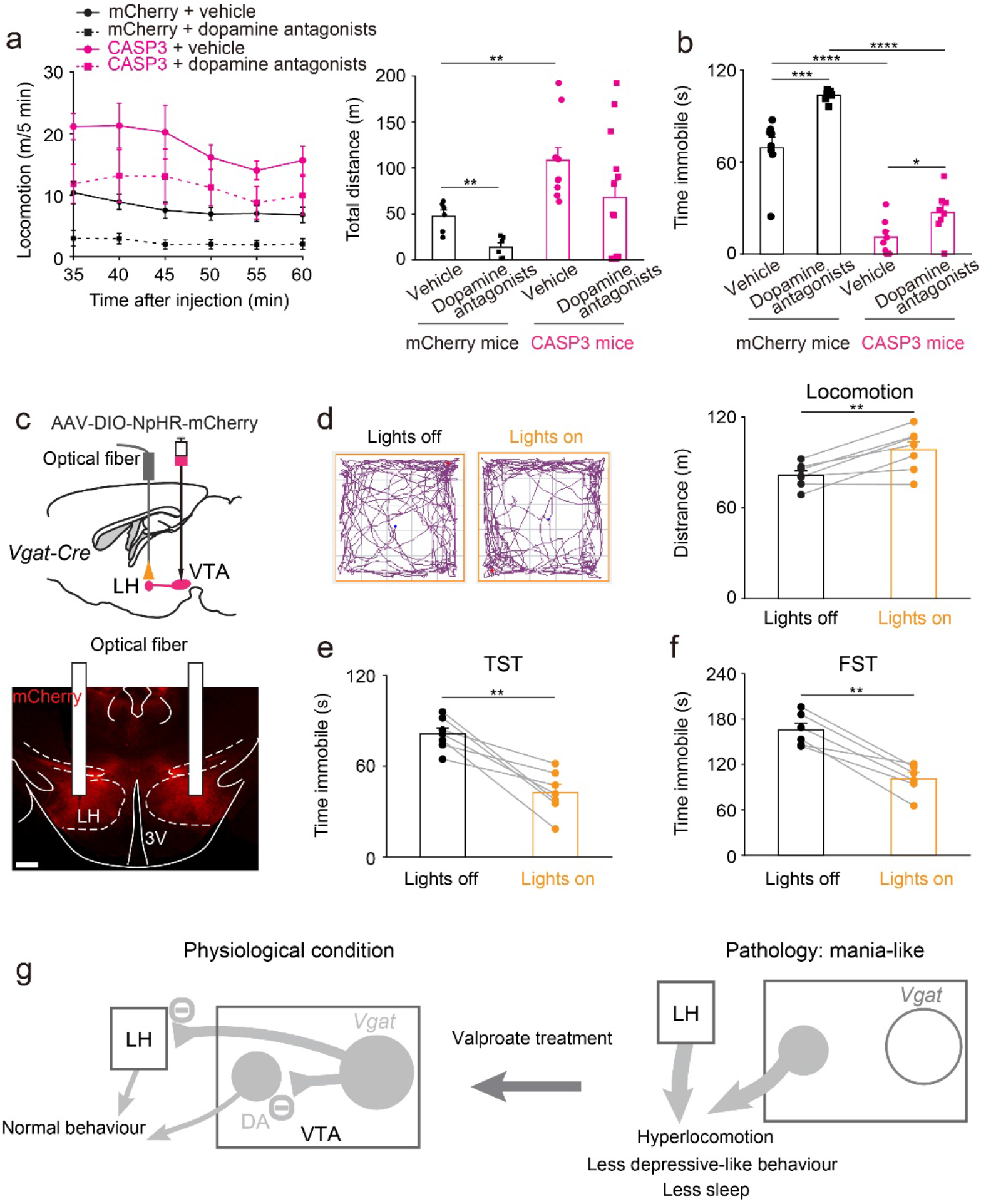
Circuit mechanisms underlying the mania-like behaviors mediated by VTA*^Vgat^* neurons. (a) Locomotion speed and distance travelled for VTA*^Vgat^*-mCherry and VTA*^Vgat^*-CASP3 mice that received injections with either vehicle (n=6 mice for mCherry and n=10 mice for CASP3) or a dopamine receptor antagonist mixture (SCH-23390 and raclopride for D1 and D2/D3 receptors, respectively) (n=6 mice for mCherry and n=13 mice for CASP3). Two-way ANOVA and Bonferroni-Holm *post hoc* test. F_(mCherry or CASP3)_=10.8; F_(saline or antagonists)_=4.5; F_(interaction)_=0.03; mCherry + vehicle vs. mCherry + antagonists t(2)=−4.14, **p=0.002; CASP3 + vehicle vs. CASP3 + antagonists t(21)=1.64, p=0.11; mCherry + vehicle vs. CASP3 + vehicle t(14)=−3.28, **p=0.005; mCherry + antagonists vs. CASP3 + antagonists t(17)=−1.91, p=0.07. All error bars represent the SEM. (b) Time spent immobile for the tail suspension assay of VTA*^Vgat^*-mCherry and VTA*^Vgat^*-CASP3 mice that received injections of either vehicle (n=8 mice for mCherry and n=9 mice for CASP3) or the dopamine receptor antagonist mixture (n=8 mice for mCherry and n=9 mice for CASP3). Two-way ANOVA and Bonferroni-Holm *post hoc* test. F_(mCherry or CASP3)_=209; F_(saline or antagonists)_=25; F_(interaction)_=4.38; mCherry + vehicle vs. mCherry + antagonists t(14)=−4.87, ***p=0.0002; CASP3 + vehicle vs. CASP3 + antagonists t(16)=−2.16, *p=0.04; mCherry + vehicle vs. CASP3 + vehicle t(15)=7.8, ****p=1e^-6^; mCherry + antagonists vs. CASP3 + antagonists t(15)=13, ****p=7e^-10^. All error bars represent the SEM. (c) AAV-DIO-NpHR-mCherry was injected the VTA of *Vgat-ires-cre* mice, and optic fibers were bilaterally implanted above the LH. The NpHR-mCherry fibers projecting from the VTA*^Vgat^* neurons into the LH were stained by immunohistochemistry (mCherry, red). Scale bar: 200 μm. (d) Video-tracked paths and distance travelled in the open field by VTA*^Vgat^*-NpHR mice (n=7 mice) with and without opto-inhibition of VTA*^Vgat^* terminals in the LH over 5 minute period. Paired t-test, t(6)=−4.08, **p=0.006. (e) Time spent immobile on the tail suspension test (TST) of VTA*^Vgat^*-NpHR mice (n=7 mice) with and without opto-inhibition of VTA*^Vgat^* terminals in the LH. Paired t-test, t(6)=5.3, **p=0.001. (f) Time spent immobile on the forced swimming test (FST) of VTA*^Vgat^*-NpHR mice (n=6 mice) with and without opto-inhibition of VTA*^Vgat^* terminals in the LH. Paired t-test, t(5)=6.77, **p=0.001. (g) Conceptual circuit diagram illustrating VTA^Vgat^ neurons inhibiting VTA dopamine (DA) neurons and circuitry in the LH. When the VTA*^Vgat^* cells have acutely diminished or absent function, activity of VTA dopamine neurons and arousal-promoting neurons in the LH increase. Valproate can reverse the effects of these changes.

VTA*^Vgat^* neurons project prominently to the LH^26^. We conducted optogenetic and chemogenetic experiments to examine if the VTA*^Vgat^* to LH projection contributes to hyperactivity and mood-related behaviors. An AAV carrying a Cre-dependent eNpHR3.0-mCherry transgene was injected into the VTA of *Vgat-ires-cre* mice to express inhibitory halorhodopsin in VTA*^Vgat^* neurons to generate VTA*^Vgat^*-NpHR mice (Figure 5c). Optic fibers were placed above the LH of VTA*^Vgat^*-NpHR mice, where dense NpHR-mCherry projections arising from the VTA*^Vgat^* cell bodies can be seen (Figure 5c). Optogenetic inhibition of the VTA*^Vgat^* to LH projection, by activating NpHR-mCherry in the LH, elevated the locomotion of mice in the open field (Figure 5d), whereas it decreased the immobility times during the TST (Figure 5e) and FST (Figure 5f). We confirmed this using specific chemogenetic inhibition of the VTA*^Vgat^* to LH projection. CNO was infused directly into the LH of VTA*^Vgat^*-hM4D_i_ mice, which expressed the inhibitory CNO receptor in VTA*^Vgat^* neurons (Supplementary Figure 5a). CNO infusion increased locomotion speed (Supplementary Figure 5b), and decreased immobility time during the TST (Supplementary Figure 5c). These results indicate that inhibiting GABAergic tone from VTA*^Vgat^* neurons to the LH produces hyperlocomotion and less depressive-like behaviors.

## Discussion

We found that endophenotypes (hyperactivity, elevated mood and reduced sleep) resembling aspects of mania could be produced by lesion or acute inhibition of GABAergic VTA*^Vgat^* neurons. In mice with lesioned or inhibited VTA*^Vgat^* neurons, D-amphetamine further increased the hyperactivity, suggesting the mania-like behavior of these mice could be the type associated with bipolar disorder in humans. The mania effects were partially blocked by D1, D2 and D3 receptor antagonists. Thus, hyperdopaminergia could originate from VTA*^Vgat^* cells failing to inhibit dopamine neurons (Fig. 5g). VTA*^Vgat^* cells, however, are not simply VTA inhibitory interneurons that reduce dopamine’s actions. Part of their actions in promoting increased movement in the FST and TST depends on their inhibitory projections to the LH. Our results on hyperactivity produced by disinhibiting LH circuits are consistent with previous findings. In rats, repeated delivery of subthreshold stimuli (kindling) to the LH induces mania-like behavior^45^. Indeed, the LH promotes physical activity and motivated behavior^46^; LH lesions in rodents and people produce a state of wakefulness with no motion^47^. The LH contains orexin/glutamate neurons that promote arousal, motivation and energy expenditure^48^, and GABAergic neurons whose activation induces wakefulness and locomotion^46, 49, 50^. It is likely that the VTA*^Vgat^* neurons inhibit both the orexin neurons and/or the arousal/locomotor-promoting GABA neurons in the LH^26^.

Inability to sleep is one of the diagnostic criteria for the mania phase of bipolar disorder^2, 3, 51^, and certainly VTA*^Vgat^* mice sleep consistently less than control mice, such that they have 100% wakefulness during the “lights off” phase compared with control mice that take about 4 hours NREM sleep during this time^26^. Usually, the longer that wakefulness persists, the stronger the urge to sleep becomes, until sleep is inescapable. A remarkable finding is that VTA*^Vgat^*-CASP3 mice bypassed the homeostatic process of having NREM recovery sleep after sleep deprivation – they did not catch up on lost sleep in spite of already starting from a strongly sleep-deprived background. The mechanisms underlying sleep homeostasis are not well understood^44, 52, 53^. Sleep homeostasis is thought to reflect the function of sleep, (otherwise why catch up on lost sleep?), and it will be interesting to investigate if the chronic lack of sleep of VTA*^Vgat^*-CASP mice will be detrimental metabolically.

Sleep deprivation can sometimes trigger mania episodes in humans^3, 51, 54^. Consequently, an interesting question is whether the chronic sleep deprivation phenotype of VTA*^Vgat^*-CASP3 mice actually causes their mania-like symptoms. Mice chronically sleep derived with the flower pot method - the animals stay on top of a raised platform surrounded by water, and when they fall asleep, they fall into water and wake up - also develop mania-like behavior^55^. This type of sleep deprivation is, however, likely to be stressful. In the VTA*^Vgat^*-CASP3 mice, the sleep-deprivation phenotype is internally generated within the brain, and could be less stressful *per se*.

The mania-like behaviors of VTA*^Vgat^*-CASP3 mice, including the strong sleep loss and abnormal sleep architecture, were reversed by valproate, and re-emerged when valproate treatment was stopped. Valproate, which is FDA-approved for treating bipolar disorder, enhances GABAergic transmission and reduces action potential firing (reviewed in ref ^13^). Lithium salts, however, had no effect in treating VTA*^Vgat^*-CASP3 mice. Lithium treatment is the first choice to treat mania episodes, although a subset of bipolar patients with rapidly cycling mania and depression phases are resistant to lithium (reviewed in ref ^13^). In most mouse models of mania, both valproate and lithium are usually effective treatments (reviewed in ref. ^1^). But similar to our results, mice with SHANK3 overexpression in the neocortex, hippocampus and basal ganglia, have mania-like symptoms treatable with valproate but not lithium^13^. It could be that the VTA*^Vgat^*-CASP3 mice and SHANK3-overexpressing mice models reflect a specific mania type.

In summary, the model based on reducing VTA*^Vgat^* neuronal function could provide further insight into the genesis of some types of mania-like behaviors. One hypothesis is that VTA*^Vgat^* neurons help set the level of mental and physical activity. Physiologically, inputs that inhibit VTA*^Vgat^* neurons will transiently intensify aspects of wakefulness useful for acute success or survival: increased activity, enhanced alertness and motivation, reduced sleep (Figure 5g). Taken to the extreme, however, pathology could emerge and decreased or failed inhibition from VTA*^Vgat^* neurons will produce mania-like qualities (Figure 5g).

## Acknowledgements

Our work was supported by the Wellcome Trust (107839/Z/15/Z, N.P.F. and 107841/Z/15/Z, W.W); the UK Dementia Research Institute (W.W. and N.P.F.); the Funds for International Cooperation and Exchange of the National Natural Science Foundation of China (Grant No. 81620108012, H.D. and N.P.F.); the China Scholarship Council (Y.M.); the research programme Rubicon (019.161LW.010) financed by the Netherlands Organization for Scientific Research (NWO) (W.B.); and the People Programme (Marie Curie Actions) of the European Union’s Eight Framework Programme H2020 under REA grant agreement 753548 (W.B.)

## Supplementary figures

**Supplementary Figure 1 (supports Figure 1):**
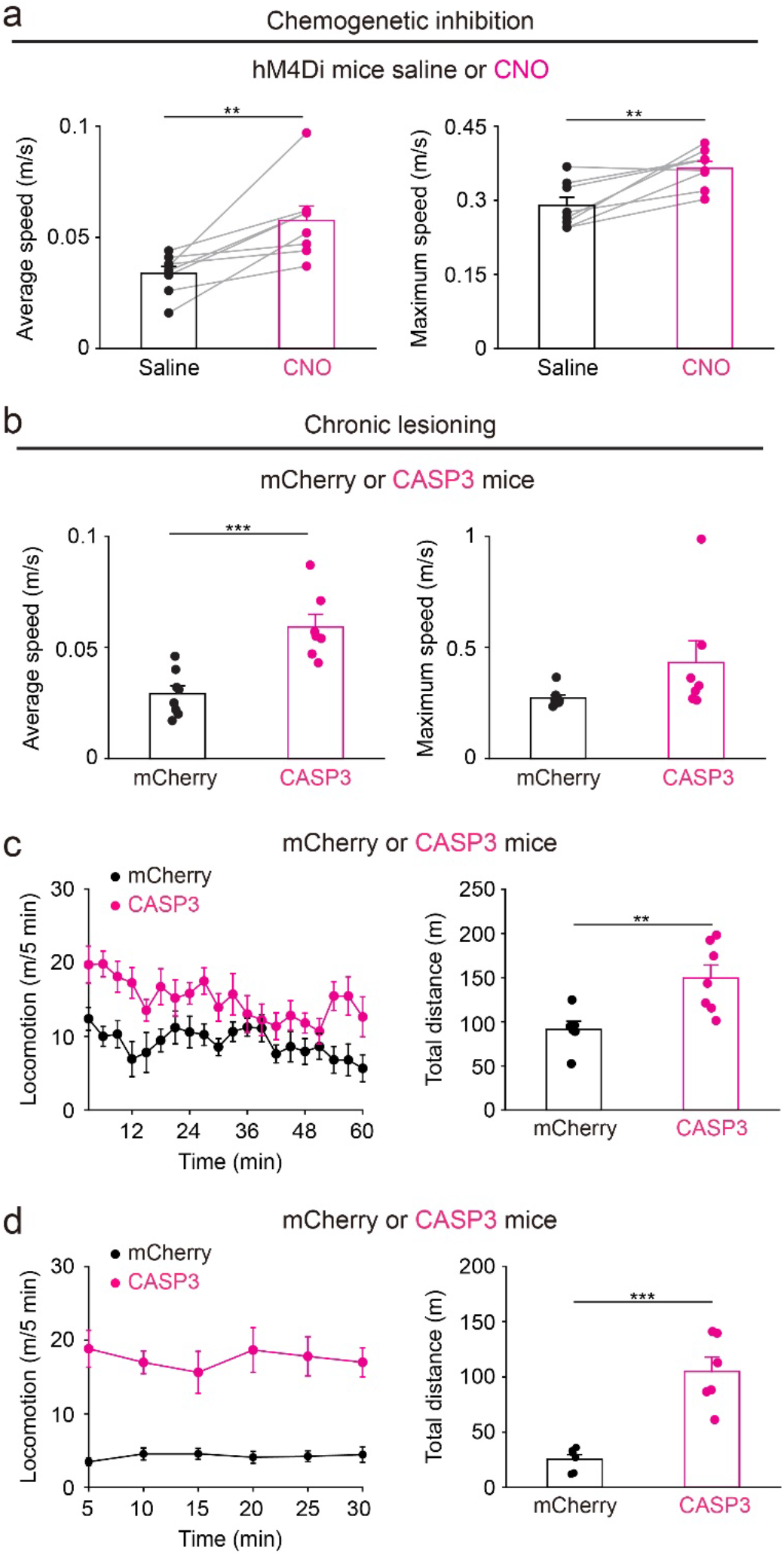
Chemogenetic inhibition or lesioning of VTA*^Vgat^* neurons produced increased locomotor activity. (a) Average speed and maximum speed of VTA*^Vgat^*-hM4Di mice (n=8 mice) that received either saline or CNO injection. Paired t-test, average speed: t(7)=−4.57, **p=0.007; maximum speed: t(7)=−3.84, **p=0.006. (b) Average speed and maximum speed of VTA*^Vgat^*-mCherry mice (n=8 mice) and control VTA*^Vgat^*-CASP3 mice (n=7 mice). Unpaired t-test, average speed: t(13)=−4.57, ***p=0.0005; maximum speed: t(13)=−1.72, p=0.1. (c) Locomotion speed and distance travelled of VTA*^Vgat^*-mCherry mice (n=6 mice) VTA*^Vgat^*-CASP3 mice (n=7 mice) in their home cages. Unpaired t-test, t(11)=−3.19, **p=0.008. (d) 4 months post-AAV injection: locomotion speed and distance travelled of VTA*^Vgat^*-mCherry mice (n=6 mice) VTA*^Vgat^*-CASP3 mice (n=6 mice). Unpaired t-test, t(10)=−5.8, ***p=0.0001.

**Supplementary Figure 2 (Supports Figure 3).**
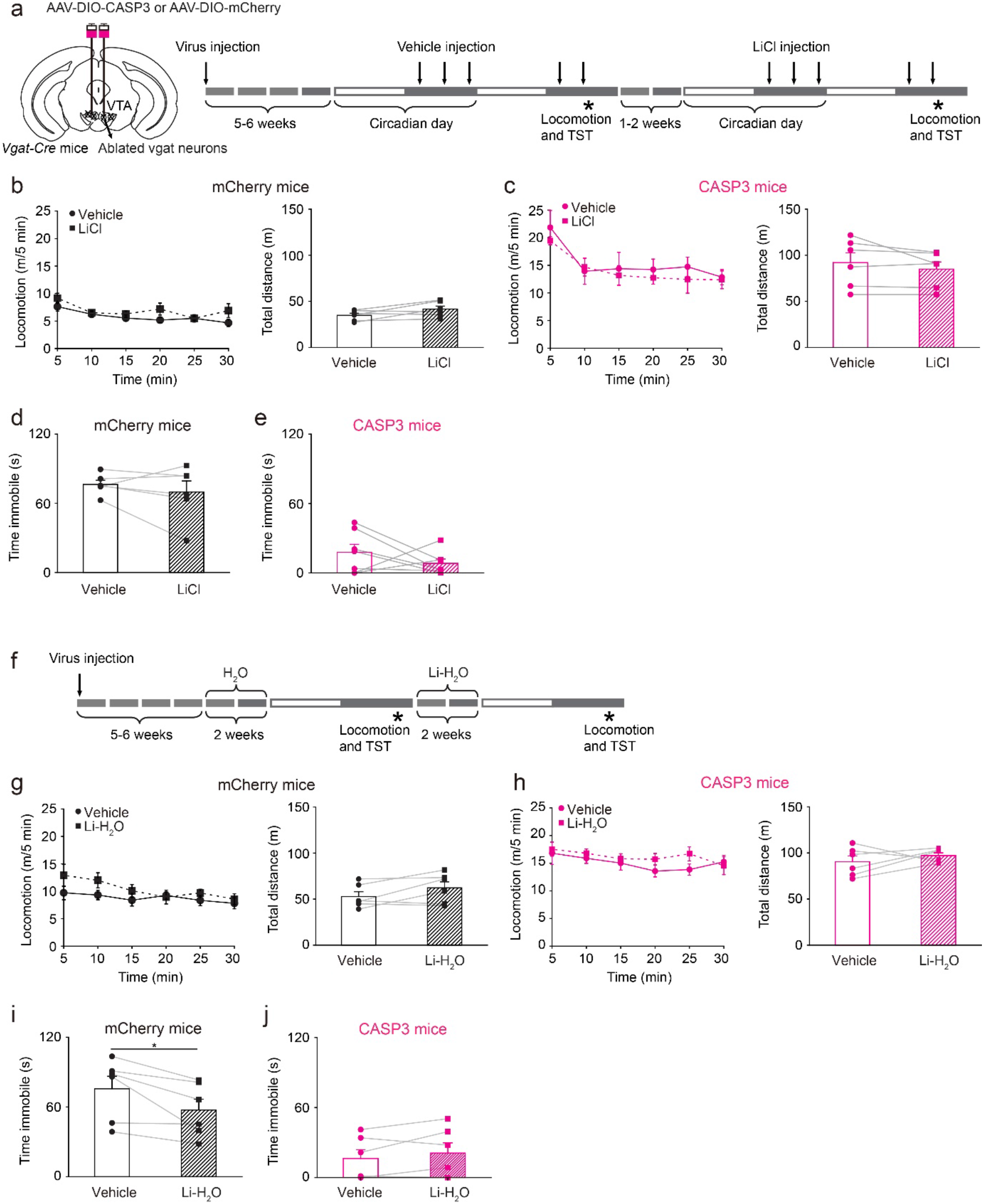
The mania-like behavior produced by lesioning VTA*^Vgat^* neurons cannot be treated by lithium. (a) Pharmacological treatment protocol for LiCl. The top arrows indicate vehicle or LiCl injection (depending on mouse group) and the stars indicate when the behavioral experiments were undertaken. (b) Locomotion speed and distance travelled of VTA*^Vgat^*-mCherry mice (n=6 mice) that received either vehicle or LiCl treatment. Paired t-test, t(5)=−2.08, p=0.09. (c) Locomotion speed and distance travelled of VTA*^Vgat^*-CASP3 mice (n=6 mice) received either vehicle or LiCl treatment. Paired t-test, t(5)=1.39, p=0.22. (d) Time spent immobile on the TST of VTA*^Vgat^*-mCherry mice (n=6 mice) that received either vehicle or LiCl injection. Paired t-test, t(5)=0.9, p=0.4. (e) Time spent immobile on the TST of VTA*^Vgat^*-CASP3 mice (n=6 mice) that received either vehicle or LiCl treatment. Paired t-test, t(5)=1.08, p=0.32. (f) Pharmacological treatment protocol for Li-H_2_O. The top arrows indicate that mice were feed with water or Li-H_2_O and the stars indicate when the behavioral experiments were undertaken. (g) Locomotion speed and distance travelled of VTA*^Vgat^*-mCherry mice (n=6 mice) that received either vehicle or Li-H_2_O treatment. Paired t-test, t(5)=−1.89, p=0.11. (h) Locomotion speed and distance travelled of VTA*^Vgat^*-CASP3 mice (n=6 mice) received either vehicle or Li-H_2_O treatment. Paired t-test, t(5)=−0.9, p=0.4. (i) Time spent immobile on the TST of VTA*^Vgat^*-mCherry mice (n=6 mice) that received either vehicle or Li-H_2_O treatment. Paired t-test, t(5)=2.85, p=0.03. (j) Time spent immobile on the TST of VTA*^Vgat^*-CASP3 mice (n=6 mice) that received either vehicle or Li-H_2_O treatment. Paired t-test, t(5)=−1.31, p=0.24.

**Supplementary figure 3 (supports Figure 3f).**
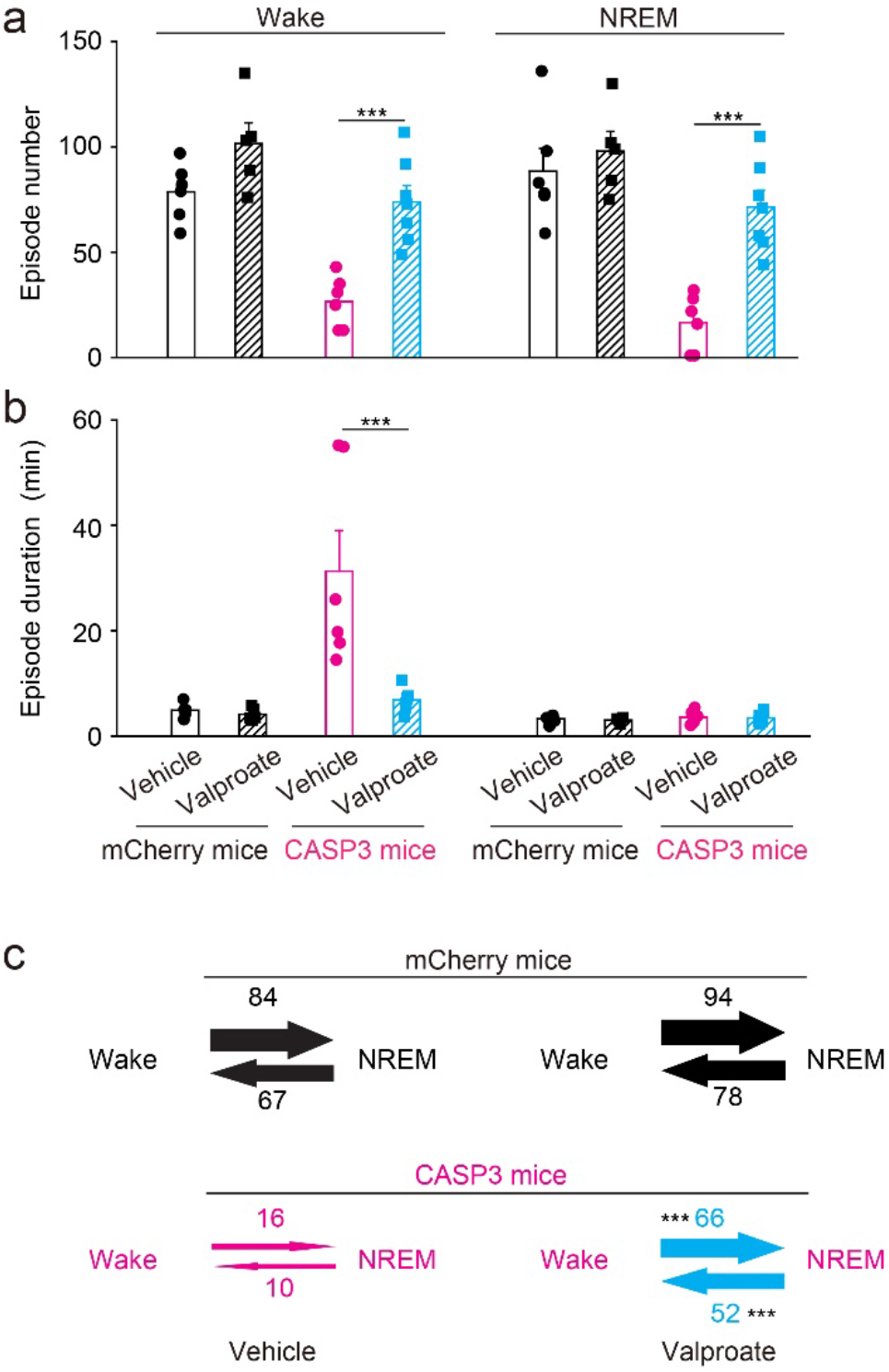
Valproate treatment normalizes the sleep-wake architecture of the VTA*^Vgat^*-CASP3 mice. (a) Episode number of wake and NREM sleep for VTA*^Vgat^*-mCherry and VTA*^Vgat^*-CASP3 mice that received vehicle or valproate injection during the 12 hours “lights off” period. mCherry + vehicle n=6 mice; mCherry + valproate n=5 mice; CASP3 + vehicle n=6 mice; CASP3 + valproate n=7 mice. Two-way ANOVA and Bonferroni-Holm *post hoc* test. Wake: F_(mCherry or CASP3)_=30; F_(Vehicle or valproate)_=24; F_(interaction)_=2.9; mCherry + vehicle vs. mCherry + valproate t(9)=−2.12, p=0.06; CASP3 + vehicle vs. CASP3 + valproate t(11)=−4.98, ***p=0.0004. NREM: F_(mCherry or CASP3)_=32; F_(vehicle or valproate)_=13; F_(interaction)_=6; mCherry + vehicle vs. mCherry + valproate t(9)=−0.64, p=0.53; CASP3 + vehicle vs. CASP3 + valproate t(11)=−5.44, ***p=0.0002. (b) Episode duration of wake and NREM sleep for VTA*^Vgat^*-mCherry mice VTA*^Vgat^*-CASP3 mice that received vehicle or valproate injection. mCherry + Vehicle n=6 mice; mCherry + valproate n=5 mice; CASP3 + vehicle n=6 mice; CASP3 + valproate n=7 mice. Two-way ANOVA and Bonferroni-Holm *post hoc* test. Wake: F_(mCherry or CASP3)_=13; F_(vehicle or_ valproate)=10; F_(interaction)_=9; mCherry + vehicle vs. mCherry + valproate t(9)=0.96, p=0.35; CASP3 + vehicle vs. CASP3 + valproate t(11)=3.45, **p=0.005. NREM: F_(mCherry or_ CASP3)=0.86; F_(vehicle or valproate)_=0.23; F_(interaction)_=0.009; mCherry + vehicle vs. mCherry + valproate t(9)=0.55, p=0.59; CASP3 + vehicle vs. CASP3 + valproate t(11)=0.24, p=0.81. (c) Transitions of wake and NREM sleep for VTA*^Vgat^*-mCherry mice and VTA*^Vgat^*-CASP3 mice that received vehicle or valproate injection. mCherry + vehicle n=6 mice; mCherry + valproate n=5 mice; CASP3 + vehicle n=6 mice; CASP3 + valproate n=7 mice. Unpaired t-test, Wake to NREM: t(11)=−5.2, ***p=0.0002; NREM to wake: t(11)=−4.46, ***=0.0009. All error bars represent the SEM.

**Supplementary figure 4 (supports Figure 5).**
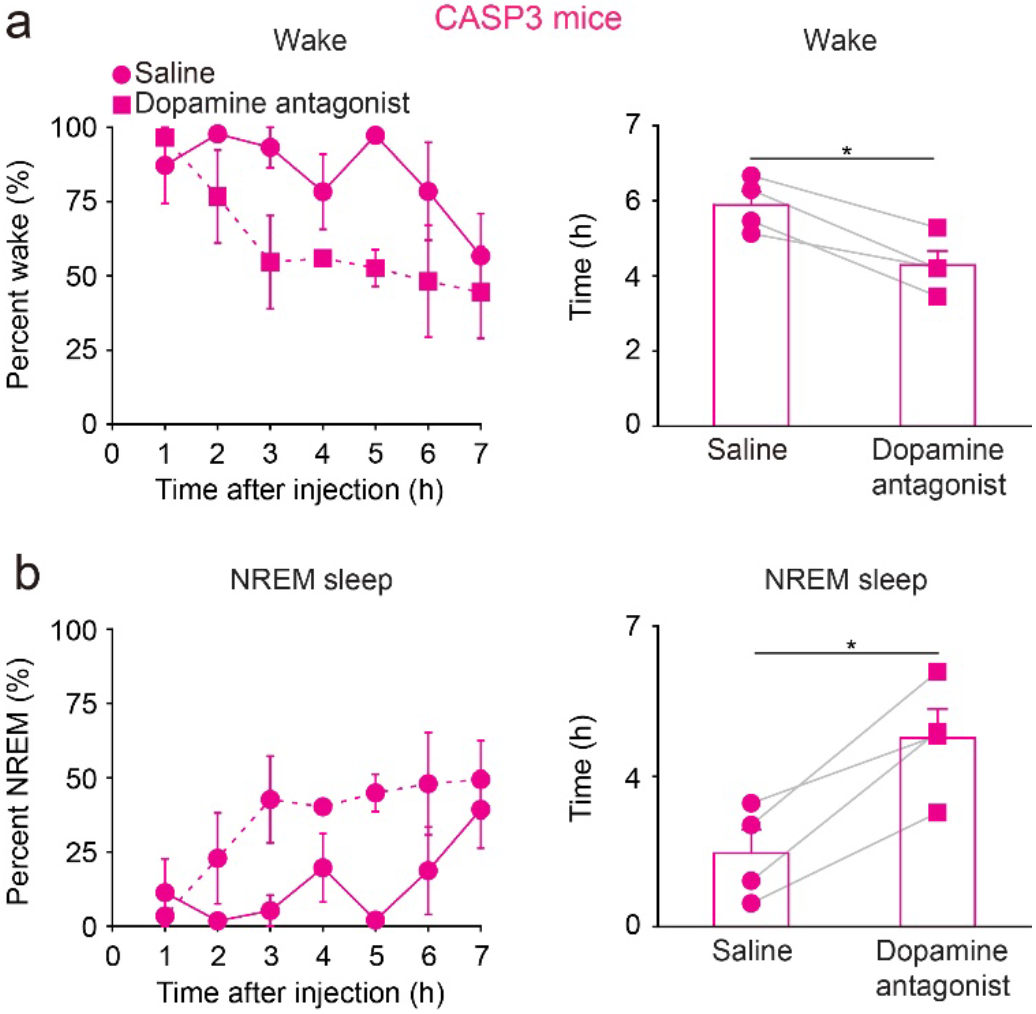
Blocking dopamine signaling restores extended wakefulness of VTA*^Vgat^*-CASP3 mice. (a) Percentage wake of VTA*^Vgat^*-CASP3 mice (n=4 mice) that received injections of saline or a dopamine receptor antagonist mixture (SCH-23390 and raclopride for D1 and D2/D3 receptors, respectively). Paired t-test, t(3)=5.75, *p=0.01. (b) Percentage NREM sleep of VTA*^Vgat^*-CASP3 mice that received either saline or the dopamine receptor antagonist mixture. Paired t-test, t(3)=−5.39, *p=0.01.

**Supplementary figure 5 (supports Figure 5).**
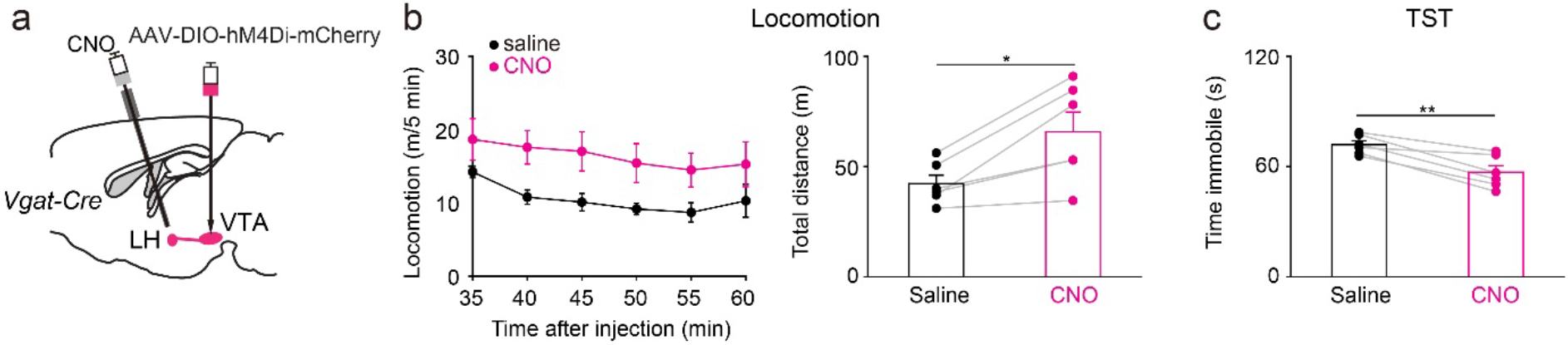
Chemogenetic inhibition of the VTA*^Vgat^* to LH projection produces hyperlocomotion and less depressive-like behavior. (a) AAV-DIO-hM4Di-mCherry was injected into the VTA of *Vgat-cre* mice. A guided-cannula to deliver CNO (5 mg/kg) or saline was implanted above the LH. (b) Locomotion speed and distance travelled for VTA*^Vgat^*-hM4Di mice (n=6 mice) that received either saline or CNO into the LH through the cannula. Paired t-test, t(5)=−3.8, *p=0.01. (c) Immobility time during the tail-suspension test (TST) for VTA*^Vgat^*-hM4Di mice (n=6 mice) that received either saline or CNO injection into the LH. Paired t-test, t(5)=5.42, **p=0.002.

